# Sporadic Diurnal Fluctuations of Cyanobacterial Populations in Oligotrophic Temperate Systems Can Prevent Accurate Characterization of Change and Risk in Aquatic Systems

**DOI:** 10.1101/2022.01.27.478008

**Authors:** Ellen S. Cameron, Anjali Krishna, Monica B. Emelko, Kirsten M. Müller

## Abstract

Continental-scale increases in aquatic system eutrophication are linked with increased cyanobacteria threats to recreational water use and drinking water resources globally. Increasing evidence suggests that diurnal vertical migration of cyanobacteria are key factors that must be considered in cyanobacterial bloom risk management. While this has been discussed in marine and eutrophic freshwater contexts, reports of diurnal vertical migration of cyanobacteria in oligotrophic freshwater lakes are scant. Typical monitoring protocols do not reflect these dynamics and frequently focus only on surface water sampling approaches, and either ignore sampling time or recommend large midday timeframes (e.g., 10AM-3PM), thereby preventing accurate characterization of cyanobacterial community dynamics. To evaluate the impact of diurnal migrations and water column stratification on cyanobacterial abundance and composition, communities were characterized in a shallow well-mixed lake interconnected to a thermally stratified lake in the Turkey Lakes Watershed (Ontario, Canada) using amplicon sequencing of the 16S rRNA gene across a multi-time point sampling series in 2018 and 2022. This work showed that cyanobacteria are present in oligotrophic lakes and their community structure varies (i) diurnally, (ii) across the depth of the water column, (iii) interannually within the same lake and (iv) between different lakes that are closely interconnected within the same watershed. It underscored the need for integrating multi-timepoint, multi-depth discrete sampling guidance into lake and reservoir monitoring programs to describe cyanobacteria community dynamics and signal change to inform risk management associated with the potential for cyanotoxin production. Ignoring variability in cyanobacterial community dynamics (such as that reported herein) and reducing sample numbers can lead to a false sense of security and missed opportunities to identify and mitigate changes in trophic status and associated risks such as toxin or taste and odor production, especially in sensitive, oligotrophic systems.

**Graphical Abstract:** 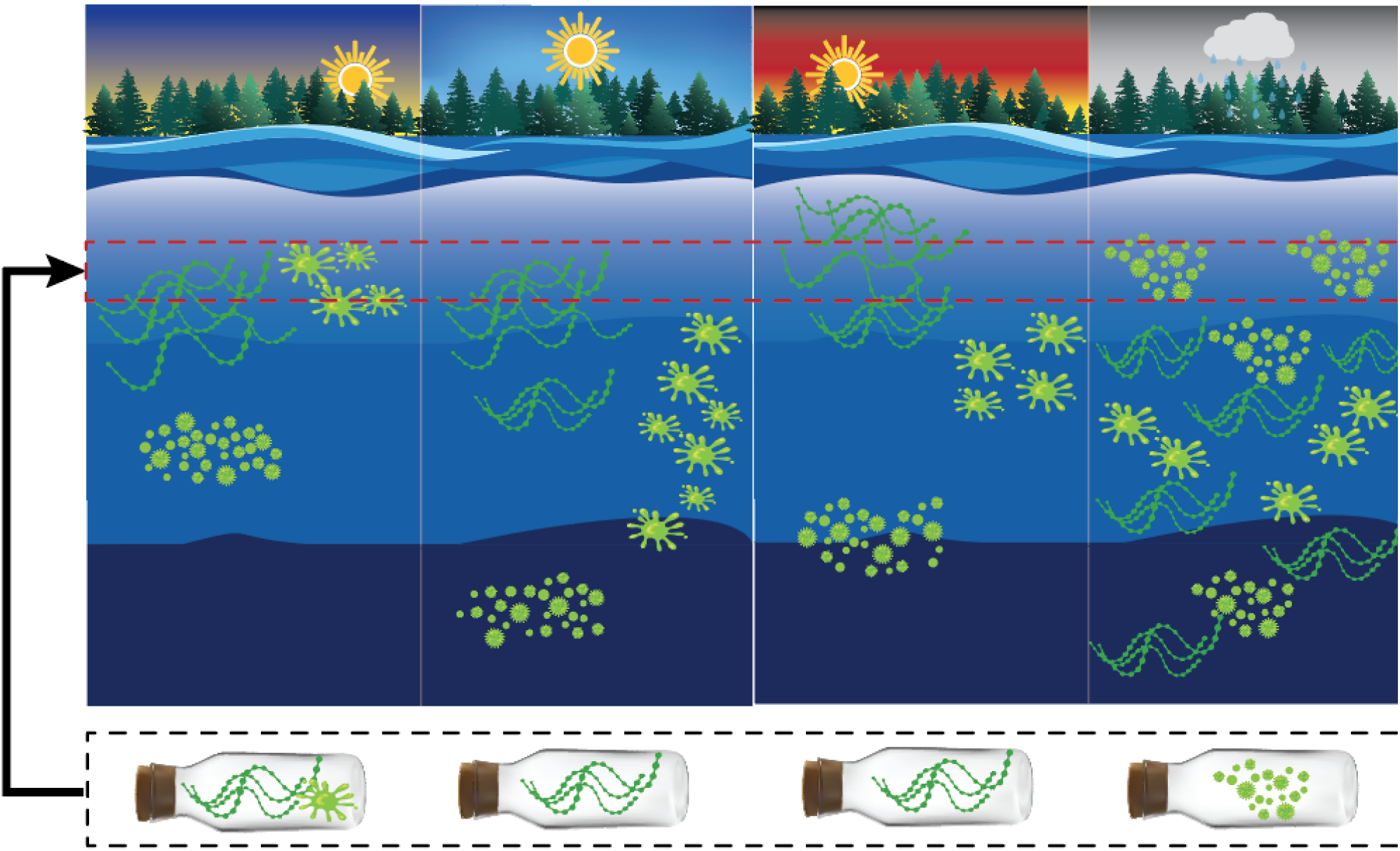

**Highlights:**

■ Cyanobacterial populations fluctuate sporadically across diurnal cycles
■ Cyanobacterial communities can vary significantly between interconnected lakes
■ Significant annual shifts in communities signal higher risk and need for monitoring
■ Cyanobacteria monitoring for risk management should incorporate time and depth

## 1. Introduction

Cyanobacteria are recognized as a threat to surface water quality as they can form dense blooms and produce secondary metabolites including potent cyanotoxins (Harke et al., 2016; Huisman et al., 2018; Paerl, 2014; Vu et al., 2020) and, compounds associated with taste and odor problems (e.g., geosmin, 2-methyl isoborneol). As a result of anthropogenic activities, cyanobacterial bloom frequency and intensity have been increasing globally (Huisman et al., 2018) and climate change-exacerbated landscape disturbances (e.g., wildfires) further promote their proliferation (Emelko et al., 2016; Silins et al., 2014). These organisms can challenge water treatment processes beyond their response capacities (Emelko et al., 2011) and exacerbate disinfection by-product formation potential after landscape disturbances such as wildfires (Tsai et al., 2017). Accordingly, continued increases in toxic bloom occurrence globally (Huisman et al., 2018) underscore the need to advance source water protection and drinking water safety plan development guidance. Improved accuracy in characterization of cyanobacterial communities and their distributions which are underpinned by diurnal migrations will accelerate that advancement and associated risk management.

The diurnal migration of cyanobacterial populations is driven by cellular characteristics (Naselli-Flores et al., 2021) and water column stability (Walsby et al., 1997), as summarized in Table S1. Diurnal migration rates (e.g., flotation and/or sinking rates) and vertical distribution of populations may be distinctive to individual taxa arising from characteristic differences in cell sizes or the presence of specialized cellular structures impacting cellular density (Naselli-Flores et al., 2021; Reynolds et al., 1987). However, these characteristic distributions are further impacted by water column stability (Walsby et al., 1997) with the potential for oscillatory diurnal variation in the distribution of cyanobacterial populations within systems of differing water column stability (Hunter et al., 2008). Rapid changes caused by oscillatory diurnal variation in the distribution of cyanobacterial populations can challenge detection if community structure and system dynamics are not well characterized.

Climate variability and landscape change-associated impacts on freshwater systems are key drivers of the need to better describe cyanobacterial growth and behavior in freshwater systems. Warming temperatures alter thermal stratification and mixing regimes in lakes (Woolway and Merchant, 2019) and increases in extreme precipitation and associated runoff increase nutrient delivery and eutrophication in receiving waters (Scholz and Brahney, 2022; Stoddard et al., 2016) and contribute to increases in cyanobacterial blooms, even in oligotrophic systems (Reinl et al., 2021; Sterner et al., 2020). Improved community characterization that reflects contemporary understanding of the diversity of cyanobacterial populations and their adaptations is therefore essential to managing risks attributable to the presence of these potentially toxic organisms in water supplies and better informing pre-emptive mitigation of potential health impacts and drinking water treatment challenges.

Key factors that must be considered in cyanobacterial bloom risk management include diurnal vertical migration of cyanobacteria, which has been reported in marine (Olli, 1999) and eutrophic freshwater (Hunter et al., 2008; von Orgies-Rutenberg et al., 2018) systems. More recently, advanced technologies including remote sensing techniques have been used to monitor cyanobacterial blooms evolution in eutrophic systems and has enabled high-resolution exploration of diurnal variability of visible bloom biomass (Li et al., 2023; Wang et al., 2023; Xue et al., 2023). In contrast, reports of diurnal vertical migration of cyanobacteria in oligotrophic freshwater lakes are scant and have been limited to spectrophotometric analysis of chlorophyll-*a* and microscopic cell enumeration over periods of 24 hours or less (Frempong, 1981; Ganf, 1974; Gilbert et al., 2010; Ibelings et al., 1991; Olli, 1999; Shahraki et al., 2020). Despite next-generation sequencing and metagenomic approaches being used to characterize cyanobacterial communities in freshwater lakes (Valadez-Cano et al., 2022) and in the context of water treatment (Jalili et al., 2021; Le et al., 2022). Studies using next-generation sequencing technology to characterize diurnal variation of cyanobacteria remain limited in number and scope, however (Gilbert et al., 2010; Shahraki et al., 2020). While the use of multi-time point sampling series provides insights into the variability observed in community composition over periods of hours, investigations that use them do not typically encompass the natural dynamic variability that aquatic systems may experience day-to-day.

Here, we provide guidance for the development of cyanobacterial monitoring programs for risk management in oligotrophic freshwater systems, the impact of sampling time and depth on cyanobacterial community composition were evaluated using multi-time point sampling over consecutive days and amplicon sequencing of the V4 region in the 16S rRNA gene. Taxonomic composition and community diversity analyses were used to examine diurnal trends in cyanobacterial distribution and the associated impact of sampling time. Fluctuations in cyanobacterial community composition were evaluated (i) temporally over a 48-hour multi-time point sampling period and (ii) spatially within the water column of a shallow well-mixed lake and an interconnected, thermally stratified lake. Temporal differences were further characterized in the thermally stratified lake during a second aquatic growing season.

## 2. Methods

### 2.1 Study Site: Turkey Lakes Watershed

The Turkey Lakes Watershed (TLW) Study was established in 1980 to investigate ecosystem effects of acidic atmospheric deposition; Jeffries et al. (1988) provide a comprehensive description of the physical characteristics of the watershed. In brief, it is approximately 50 km north of Sault Ste Marie, Ontario on the Canadian Shield in an uneven-aged tolerant hardwood and mixed conifer forest (Jeffries et al., 1988). This watershed consists of four interconnected lakes fed by both first order streams and groundwater: Batchawana Lake, Wishart Lake, Little Turkey Lake, and Big Turkey Lake (Jeffries et al., 1988; Figure S1). The shallow depth of Wishart Lake often results in complete wind-induced mixing that prevents water column stratification whereas Little Turkey Lake, Big Turkey Lake and Batchawana Lake are deeper and regularly undergo thermal stratification (Figure S2). These lakes are classified as oligotrophic to mesotrophic, and cyanobacteria have been reported as the dominant members of phytoplankton communities and previously reported as major contributors to peaks in primary production during summer periods (Jeffries et al., 1988).

### 2.2 Sample Collection

Water samples were collected from the deepest point in Little Turkey (47°02’37.2”N 84°24’24.4”W) and Wishart (47°03’00.0”N 84°23’58.3”W) Lakes during a two-day period across three time points on both days (2018a/b, 2022a/b): morning (8-9 a.m.), midday (12-1 p.m.) and afternoon (4-5 p.m.) during two sampling events (August 2018 and August 2022). Samples were collected from Secchi depth, the water depth at which light penetration is approximately 1% of surface illumination and is considered the maximum depth at which there is sufficient light for photosynthesis (Bukata et al., 1988). In Wishart Lake, these samples were collected from near the bottom (4 m) due to high water clarity. Secchi depths ranged from 4 to 5.25 m in Little Turkey Lake. Water samples were also collected at the surface (0 m) in Little Turkey Lake on the first sampling day in 2018 (2018a) and both sampling days in 2022 (2022a, 2022b) to evaluate potential temporal correlations in cyanobacterial communities between depths.

Water samples collected using a Masterflex E/S portable sampler peristaltic pump were vacuum filtered through a 47 mm GF/C filter (Whatman plc, Buckinghamshire, United Kingdom). For samples collected in August 2018, 250 mL of vacuum filtered water were filtered a second time through a 0.22 µm Sterivex™ filter to collect additional microbes. All filters were stored at -20°C prior to DNA extraction. Sampling details are provided in Table S2.

### 2.3 DNA Extraction, V4 16S rRNA Gene Amplicon Sequencing

DNA was extracted using the DNeasy PowerSoil Kit (QIAGEN Inc., Venlo, Netherlands) following the manufacturer’s protocol. Following this, elution buffer was added to spin columns for 15 minutes prior to elution. DNA was quantified using a NanoDrop spectrophotometer (Table S2); absolute values were only accurate at DNA concentrations of more than 10ng/µl. The DNA extracts were submitted to a commercial laboratory (Metagenom Bio Inc.,Waterloo, ON) for amplicon sequencing. Primers designed to target the 16S rRNA gene V4 region [515FB (GTGYCAGCMGCCGCGGTAA) and 806RB (GGACTACNVGGGTWTCTAAT)] (Walters et al., 2015) were used for PCR amplification. Polymerase chain reactions (PCR) were conducted in triplicate for each sample (25 μL). Reaction mixtures were composed of 2.5 μL of 10X *Taq* buffer, 0.5μL of 10mM dNTP, 0.25 μL of BSA (20mg/mL), 5.0 μL of 1 μM forward primer, 5.0 μL of 1 μM of reverse primer, 5.0 μL of sample DNA, 0.2 μL *Taq* DNA polymerase (5u/μL) and 6.55 μL of PCR water. DNA was initially denatured at 95°C for 5 minutes, followed by 35 cycles of: 1) 95°C for 30 seconds, 2) 30°C for 30 seconds, 3) 72°C for 50 seconds. A final extension was performed at 72°C for 10 minutes. Triplicate PCR products were pooled, gel purified and quantified using Qubit dsDNA HS Assay Kit (Thermo Fisher Scientific Inc.). Library DNA was prepared and sequenced with MiSeq Reagent Kit v2 (2x 250 cycles) and sequenced with the Illumina MiSeq platform (Illumina Inc., San Diego, United States).

### 2.4 Sequence Processing & Library Size Normalization

The program QIIME2 (v. 2019.10; Bolyen et al., 2019) was used for bioinformatic processing. Demultiplexed paired-end sequences were trimmed and denoised, including the removal of chimeric sequences and singleton sequence variants, using *DADA2* (Callahan et al., 2016). Amplicon sequence variants across the sample sets were clustered at 99% identity to account for single nucleotide discrepancies in sequenced amplicons using *vsearch* (Rognes et al., 2016) to generate a representative set of operational taxonomic units (OTUs). Taxonomic classification was performed using a Naïve-Bayes taxonomic classifier trained using the SILVA138 database (Quast et al., 2013; Yilmaz et al., 2014). Sequences classified as cyanobacteria at the phylum level were aligned to reference 16S rRNA cyanobacterial sequences in the Cyanoseq database (Lefler et al., 2023) using *MAFFT* (v7.490; Katoh and Standley, 2013). A phylogenetic tree was generated using *iqtree* (v. 2.2.0; Nguyen et al., 2015) using an unrestricted model (UNREST+FO+R10) selected by the ModelFinder Bayesian Information Criterion and visualized with Interactive Tree of Life (iTOL; Letunic and Bork, 2021; Figure S3). Confirmed taxonomic classifications for OTUs classified as cyanobacteria at the phylum level were manually curated to reflect taxonomic levels above the genus level according to taxonomic labels in the Cyanoseq database based on placements in the phylogenetic tree.

Files from QIIME2 were imported into R (v. 4.0.1; R Core Team, 2020) for downstream analyses using *qiime2R* (v. 0.99.23; Bisanz, 2018). Initial sequence libraries were filtered to exclude OTUs that were taxonomically classified as mitochondria or chloroplast sequences using *phyloseq* (v. 1.32.0; McMurdie and Holmes, 2012). For cyanobacterial community analysis, OTUs classified as cyanobacteria at the phylum level were selected to create libraries consisting of only cyanobacterial sequences.

### 2.5 Cyanobacterial Communities - Taxonomic Composition & Diversity Analyses

Bacterial community composition was assessed at the phylum level and relative abundances were visualized using a stacked bar plot produced with *mirlyn* (Cameron and Tremblay, 2020). Relative abundances were randomized across bacterial phyla within samples to identify significantly dominant groups within the bacterial community in relation to sampling time and adjusted with a Bonferroni correction using the *randomseqsig* function in *mirlyn*. Cyanobacterial community composition was assessed through the selection of a subset of sequences classified as “cyanobacteria” at the phylum level and relative abundances of cyanobacterial families were also visualized using a stacked bar plot. For bacterial and cyanobacterial community composition bar plots, groups with an average relative abundance lower than 1% across samples were collapsed into a low abundance category for visualization purposes.

As amplicon sequencing only indirectly and partially represents source diversity and clustering, amplification, and the lack of certainty about how many zeros should be included in the data compromise statistical inference about source diversity (Schmidt et al., 2021), community diversity analyses were performed using relative abundances instead of rarefying. While rarefying can be utilized to compare libraries in a statistically meaningful manner (Cameron et al., 2021), library sizes between samples collected in 2018 and 2022 differed by orders of magnitude creating extreme data loss of sequences or entire samples. OTUs present at >1% in the bacterial community were utilized for diversity analyses to reduce the risk of accidental non-detection. Alpha diversity was assessed using direct counts of unique OTUs and the Shannon Index (Shannon, 1948) to identify trends in sample diversity as a function of time. Bray-Curtis dissimilarity metric was calculated on compositional community data using *vegdist* in *vegan* (Oksanen et al., 2022) and used for permutational analysis of variance with *adonis2*. Bray-Curtis distances were visualized using ComplexHeatMap (Gu, 2022). To identify individual OTUs driving compositional changes, highly abundant (>10% of cyanobacterial community) were selected and normalized using Z-scores to identify enrichment or under-enrichment of individual abundant OTUs as a function of time.

### 2.6 Data Availability

Sequence data analyzed in this study is available in the European Nucleotide Archive (ENA) under study accession ERP134980.

## 3. Results

### 3.1 Diurnal Conservation of Bacterial & Cyanobacterial Communities

Cyanobacteria classified sequences were found to comprise a variable proportion of the bacterial community composition across sampling events in 2018 and 2022 (Figure 1A). Relative abundances of cyanobacteria were more similar during sampling events in Little Turkey Lake than Wishart Lake in 2018 (Figure 1A; Figure 2A). Cyanobacteria did not contribute to a significant portion of the bacterial community in all sampling events of 2018 (p = 1; Table S4). In contrast, cyanobacterial sequences comprised a significant portion of the bacterial community in Little Turkey Lake in 2022. Specifically, cyanobacteria comprised a significant portion of the bacterial community (i) at all sampling times at Secchi depth (p < 0.005) on both sampling days (2022a; 2022b) and (ii) in morning and midday surface samples on the second sampling day (2022b; p < 0.003).

**Figure 1:**
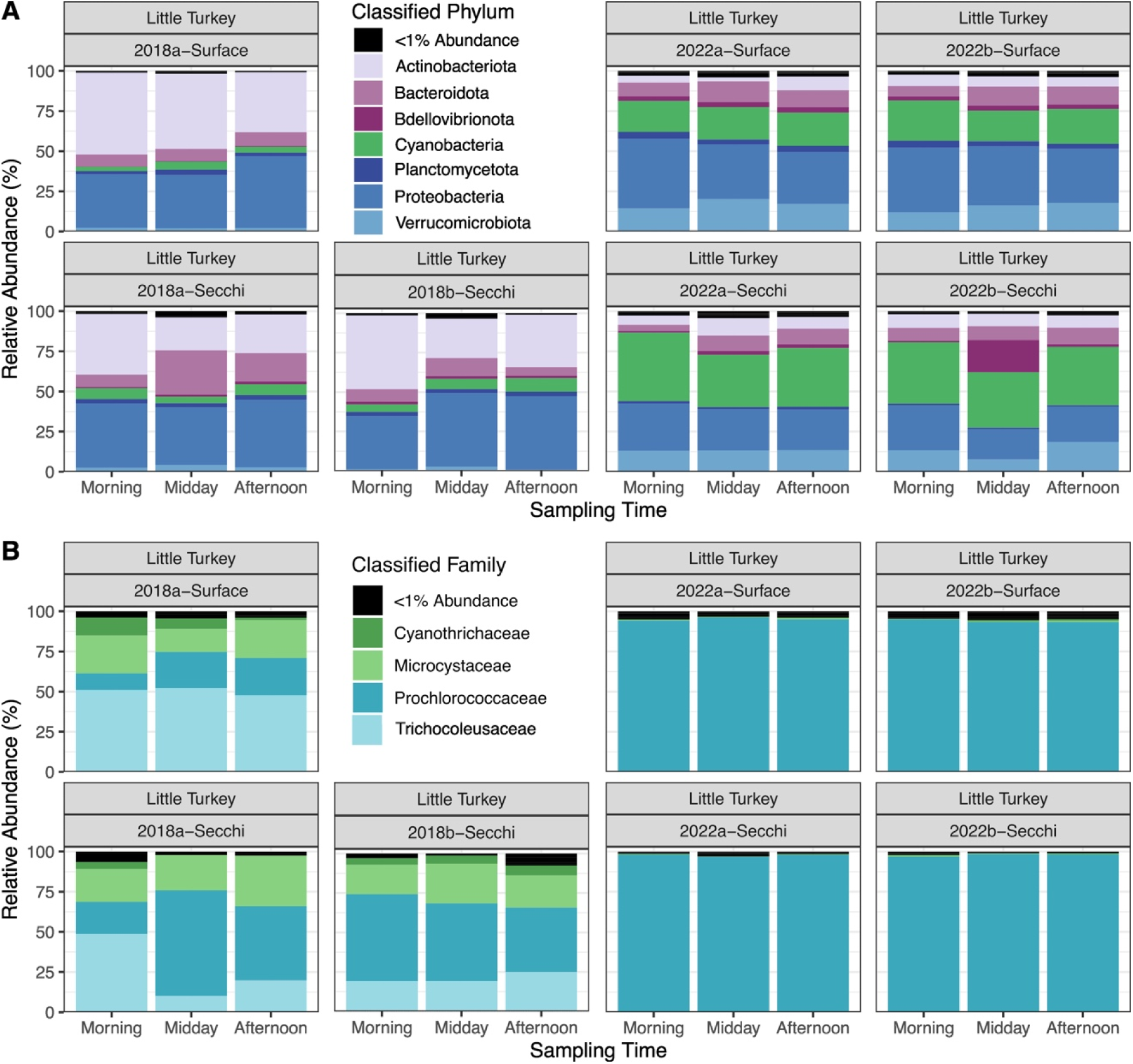
Stacked bar plots depicting the taxonomic composition of bacterial communities in terms of the relative abundance of major bacterial phyla (A) and cyanobacterial communities in terms of relative abundance of major cyanobacterial families (B) measured across a multi-time point sampling series over two consecutive sampling days (2018a/2018b, 2022a/2022b) in a thermally stratified lake (Little Turkey Lake).

**Figure 2:**
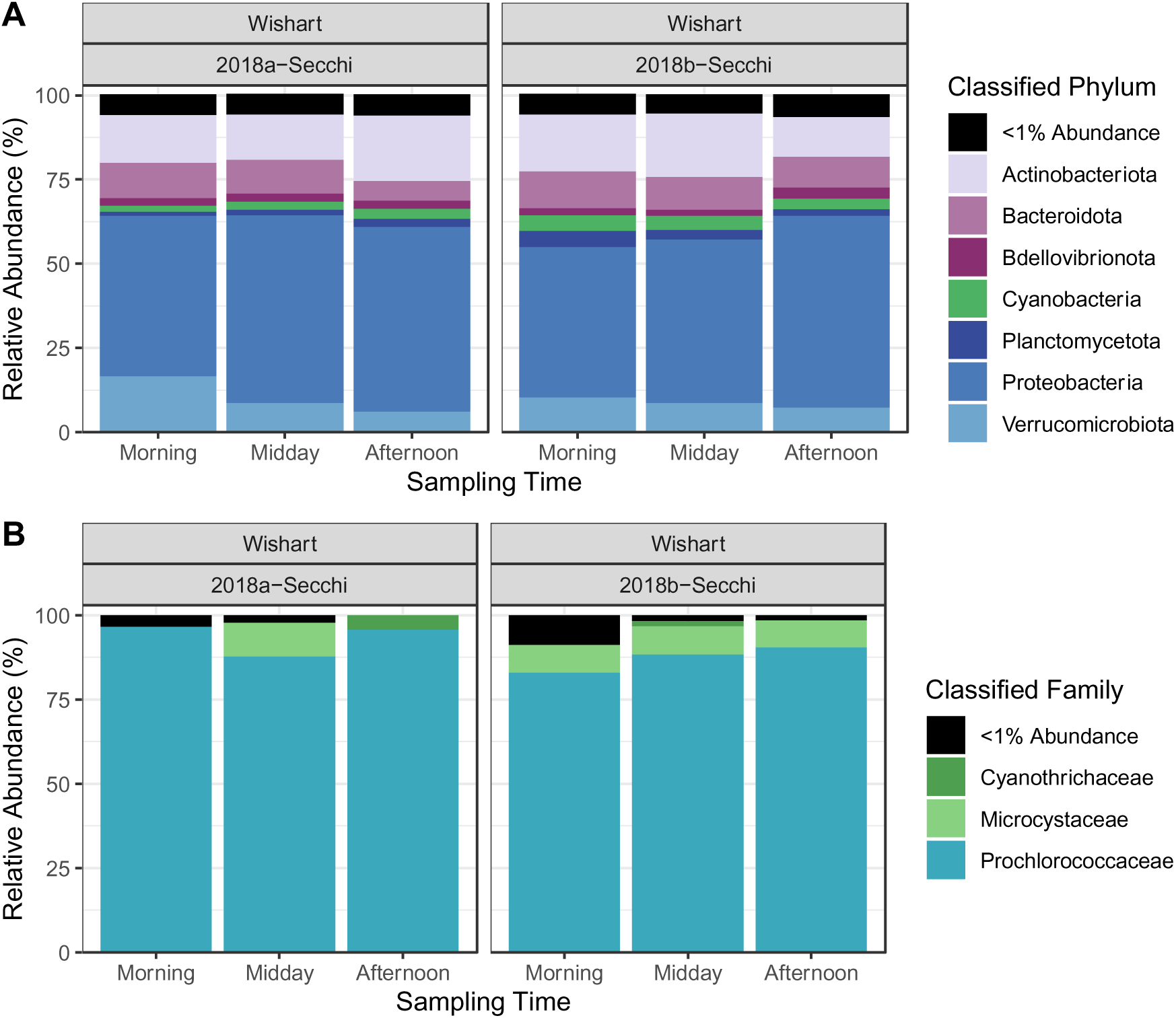
Stacked bar plots depicting the taxonomic composition of bacterial communities in terms of the relative abundance of major bacterial phyla (A) and cyanobacterial communities in terms of relative abundance of major cyanobacterial families (B) measured across a multi-time point sampling series over two consecutive sampling days (2018a/2018b) in a shallow, non-stratified lake (Wishart Lake).

Cyanobacterial communities were frequently dominated by sequences classified to the family Prochlorococcaceae with minimal diurnal fluctuations (Figure 1B; Figure 2B). Cyanobacterial communities observed in Little Turkey Lake in 2018 included large proportions of taxonomic families not observed in Wishart Lake or during the 2022 sampling events. A higher abundance of Cyanothrichaceae, Microcystaceae, and Trichocoleusaceae classified sequences were observed in the 2018 samples from Little Turkey Lake. Relative abundances of Cyanothrichaceae decreased from morning (11.1%) to afternoon (1.4%) at surface depths, possibly due to downward vertical migration. At Secchi depth on the first sampling day (2018a), Trichocoleusaceae were found in similar relative abundances (48.6%) in the morning sampling event but experienced a drastic decrease by midday (10.1%) before a second increase at the final sampling time (19.8%). On the second sampling day (2018b) in Little Turkey Lake, Trichocoleusaceae relative abundances were relatively consistent from morning (19.0%) to midday (19.1%) and then increased in the afternoon (25%). Notably, the relative abundances observed in the morning sampling point at Secchi depth of the 2018a sampling event had higher similarity to those at the water surface than those at Secchi depth during subsequent sampling events. Fluctuations in Prochlorococcaceae relative abundances were negatively correlated with Trichocoleusaceae and Microcystaceae abundances due to the compositional constraint of amplicon sequencing data.

Similarity between cyanobacterial communities across sampling events was analyzed using the Bray-Curtis dissimilarity metric (Figure 3). Sampling time did not significantly impact cyanobacterial community composition in either of the study lakes (p = 0.97; Table S5) suggesting diurnal conservation of cyanobacterial community composition. However, significant compositional differences were observed between the two lakes sites (p = 0.002). The insignificance of sampling time and significance of sampling site were also observed for bacterial community composition (p = 0.99; p = 0.001; Table S6).

**Figure 3:**
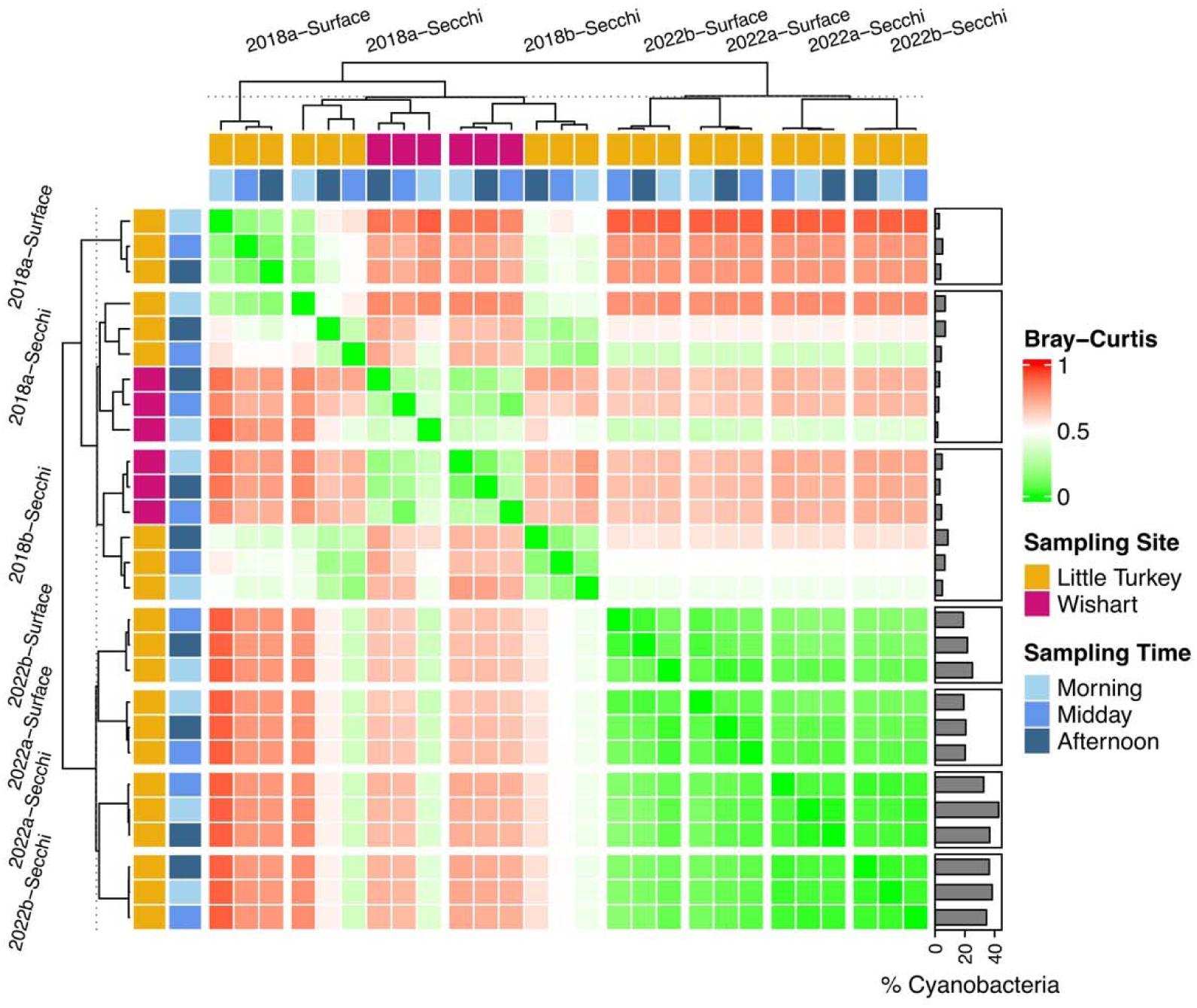
Compositional similarity of cyanobacterial communities across a multi-time point sampling series conducted over two consecutive sampling days (2018a/2018b, 2022a/2022b). Bray-Curtis dissimilarity metric (0 = completely identical, 1 = completely dissimilar) characterizes the differences present in the OTU composition of cyanobacterial communities. Overall proportion of cyanobacteria in bacterial communities displayed on the right-hand side highlights and contextualizes the interannual variability observed within Little Turkey Lake bacterial communities.

**Figure 3:**
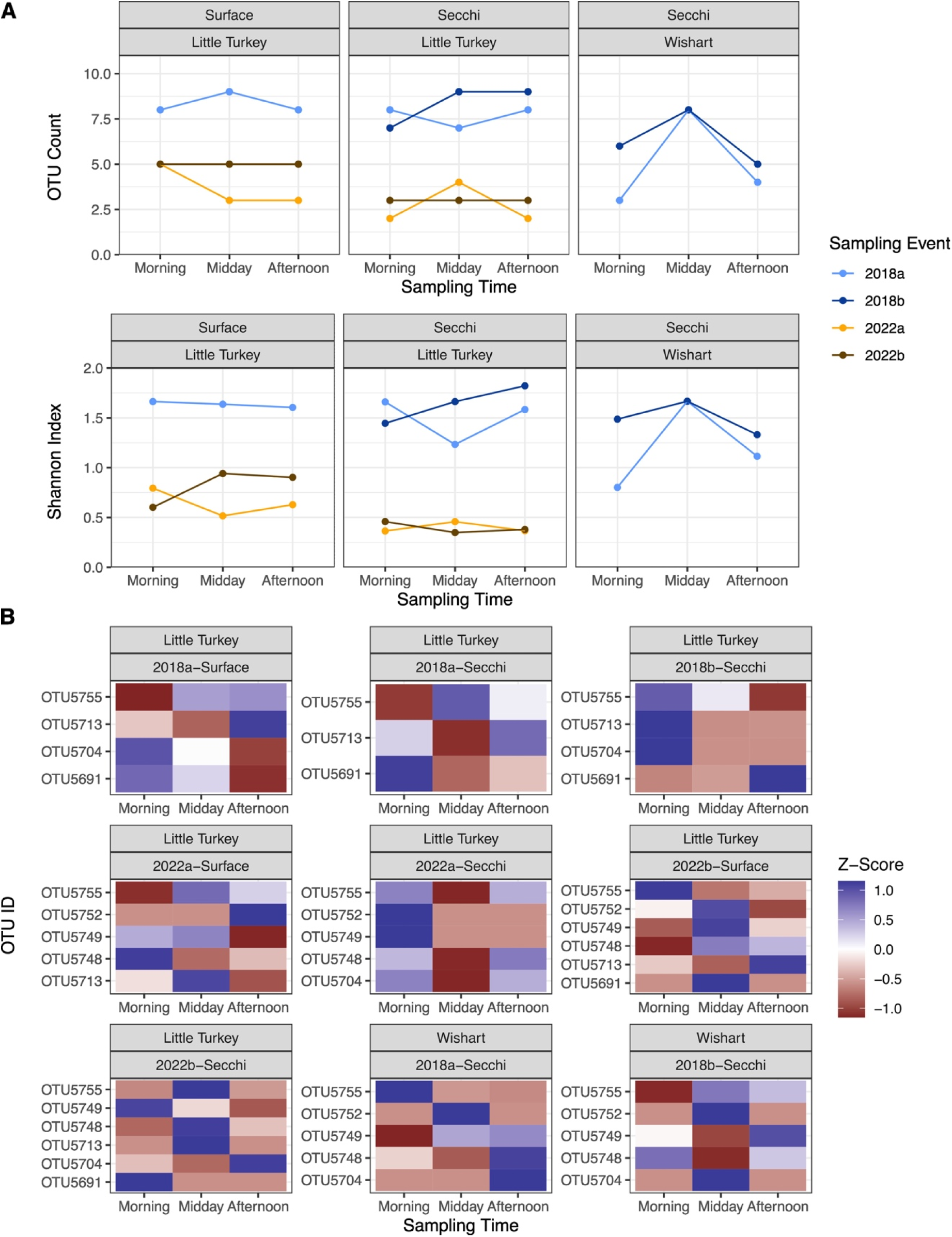
Temporal Fluctuations in alpha-diversity metrics of cyanobacterial communities driven by changes in OTU detection (OTU counts; A) and composition (Shannon Index; B). Z-score normalized relative abundances for highly abundant operational taxonomic units (>10% of cyanobacterial community) for sampling event to demonstrate diurnal trends of specific cyanobacterial populations (C).

Within Little Turkey Lake, interannual variability between sampling events had a significant impact on cyanobacterial communities (p = 0.001). Sampling depth of the stratified water column in Little Turkey did not significantly impact the community structure (p = 0.154) and high similarity was still observed between samples collected at the same sampling depth. Additionally, similarity was observed within lakes across the sampling days of each year, except in the unique case of the morning of the first sampling day of 2018 (2018a). Cyanobacterial communities at Secchi depth in the morning of the 2018a sampling event were highly similar to surface communities observed in Little Turkey Lake, a phenomenon that was not observed during other sampling events (Figure 3).

### 3.2 Sporadic Diurnal Variability in Relative Abundances of OTUs

Cyanobacterial community diversity across sampling times was assessed using (i) counts of unique OTUs (>1% abundance) and (ii) the Shannon Index (Figure 3A). Little Turkey Lake exhibited different trends in sample diversity dependent on the sampling event. Within samples collected during the 2018 sampling events, a higher number of cyanobacterial OTUs were detected in comparison to the 2022 sampling series. However, there was no consistent trend of diurnal fluctuations, revealing the sporadic fluctuations which may occur throughout a diurnal period. For example, 2018 sampling events at Secchi depth revealed opposing trends at midday where diversity scores decrease during the 2018a sampling event but increase during the 2018b sampling event. Similar results are also observed during the 2022 sampling events. The 2022b sampling event also reveals a consistent OTU count across sampling times, but the Shannon Index suggests changes in proportionality of these detected OTUs across time. Although results from Wishart Lake are limited to the 2018 sampling events, contrary to the sporadic diurnal fluctuations of diversity observed in Little Turkey Lake, Wishart Lake was shown to have corresponding spikes in diversity at the midday sampling point on both sampling days.

Diurnal fluctuations of highly abundant cyanobacterial OTUs (>10%) were identified using Z-scores; time points associated with a higher Z-score reflect greater deviation from the mean (Figure 3B). Additionally, temporal distribution of cyanobacterial OTUs was visualized to identify ubiquitous OTUs within cyanobacterial communities (Figure 4). Specifically, OTU5755 (Prochlorococcaceae) was found in all samples independent of sampling site, sampling year or sampling time. Diurnal fluctuations in the occurrence of OTU5755 could not be discerned based on the Z scores; both enrichment and de-enrichment were observed. Absence of diurnal linked fluctuations was also observed for other highly independent OTUs including those from families which may be associated with gas vacuolate taxa (e.g., Trichocoleusaceae – OTU5691). Distribution of OTUs across sampling events indicated that the predominant populations varied temporally (Figure 4). While OTU5691 and OTU5695, both classified to the family Trichocoleusaceae and comprised a significant portion of cyanobacterial communities in 2018, they were rarely detected and detected at substantially lower relative abundances in the 2022 sampling events. 2022 sampling events were found to have a higher count of unique Prochlorococcaceae OTUs than those detected in 2018.

**Figure 4:**
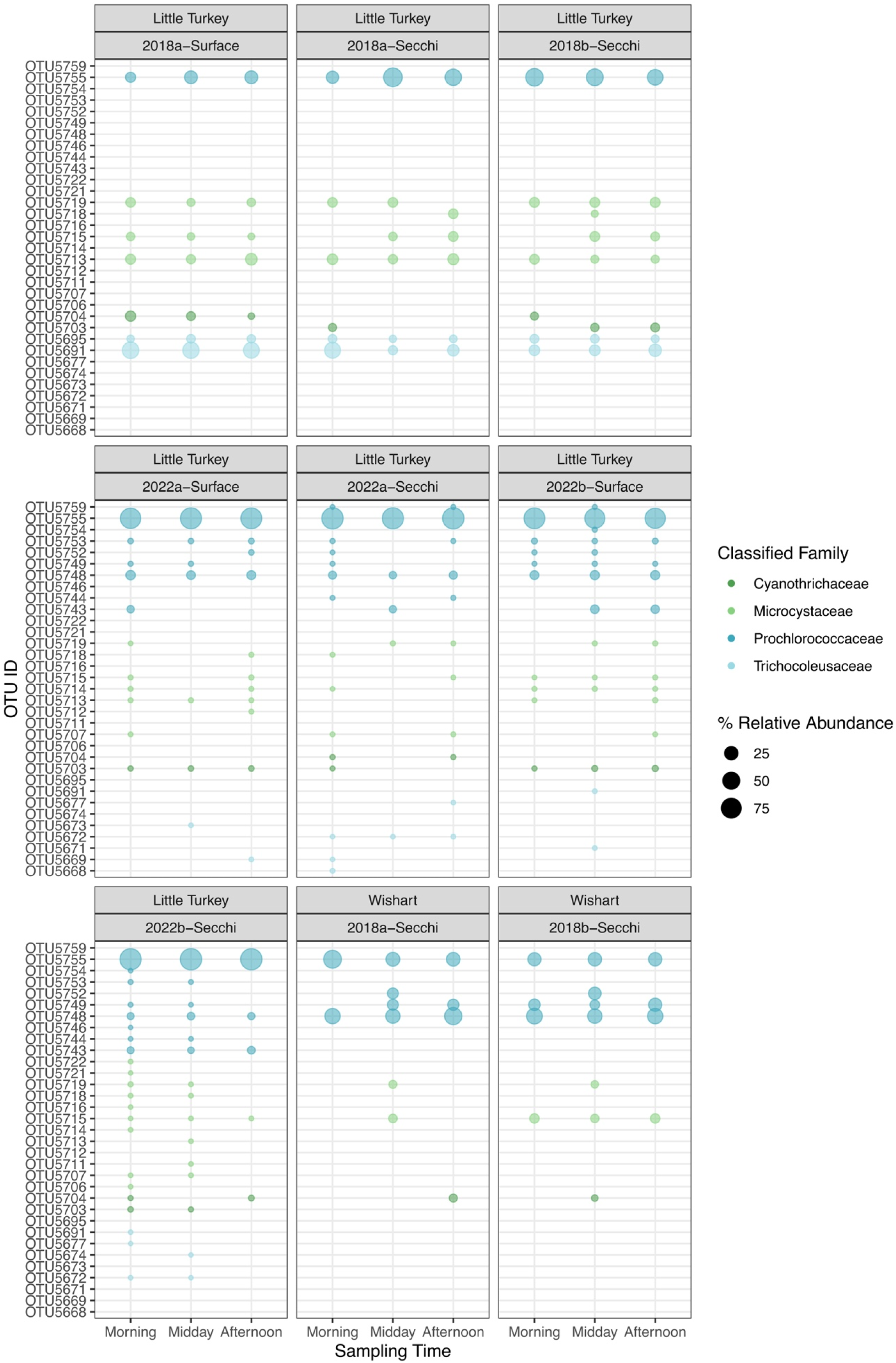
Temporal distribution of individual OTUs of cyanobacterial families present in >1% average relative abundances coloured based on family level taxonomic classification. Sizes of points represent relative abundance of OTU in cyanobacterial community.

## 4. Discussion

Diurnal migrations of cyanobacterial populations have been observed in marine and eutrophic freshwater systems and attributed to both cellular characteristics (Naselli-Flores et al., 2021) and water column stability (Walsby et al., 1997). Oscillatory diurnal variation in the distribution of cyanobacterial populations would thus be expected across depth in the water column. In contrast to trends frequently observed in marine and eutrophic freshwater systems, we demonstrate that in highly oligotrophic systems, diurnal fluctuations of cyanobacterial populations are sporadic, but overall bacterial community composition is conserved temporally. These observations contradict the patterns commonly described for diurnal migrations of gas vacuolate taxa. Consequently, these contrasting results—especially the unpredictable nature of cyanobacterial population distributions—have significant implications for the development of water quality monitoring programs and associated risk management strategies, including source water protection and drinking water safety plans.

Evidence of diurnal migrations of colonial and filamentous cyanobacteria has been reported previously; larger colonies have been shown to migrate across greater depths more rapidly than smaller ones (Reynolds et al., 1987; Westwood and Ganf, 2004); they also experience faster sinking rates (Ganf, 1974). In both Little Turkey and Wishart Lakes, cyanobacterial communities were predominantly composed of OTUs classified to the family Prochlorococcaceae. Notably, the family Prochlorococcaceae includes small unicellular and colonial taxa that may facilitate their ability to maintain water column position (Śliwińska-Wilczewska et al., 2018). Furthermore, smaller colonies also experience more random spatial movement (Chien et al., 2013). The dominance of small unicellular and colonial taxa of the Prochlorococcaceae family concurrent with the absence of visible cyanobacterial biomass may thus help to explain the absence of discernible diurnal trends in the studied cyanobacterial OTUs.

Previous studies have demonstrated that thermal stratification of the water column plays a significant role in shaping the distribution of cyanobacteria and other phytoplankton through the creation of discrete zones in the water column (*i.e.*, nutrient depleted, light rich surface waters *vs.* nutrient rich, light limited deep waters; Reynolds et al., 1987; Spigel and Imberger, 1987). To investigate the impact of thermal stratification on cyanobacterial communities in an oligotrophic lake (*i.e*., Little Turkey Lake), samples were collected in both the epilimnion (water surface) and metalimnion (Secchi depth) zones (Figure S2). During both sampling events this analysis revealed common OTUs over time and across depth, thereby suggesting a broader distribution of cyanobacterial species across the water column rather than restriction to specific zones. In 2022, the composition of the cyanobacterial communities was highly similar regardless of spatio-temporal differences. In contrast, in 2018, cyanobacterial communities at the water surface and Secchi depth were distinguishable, consistent with previous spatial distribution research conducted in Little Turkey Lake (Cameron et al., 2022). On both occasions, sampling time did not significantly impact the cyanobacterial community composition, further supporting the hypothesis that small unicellular and colonial cyanobacterial populations will exhibit random fluctuations.

Notably, cyanobacterial communities of Little Turkey Lake in 2018 and 2022 did not exhibit similarity. This result is consistent with seasonal studies conducted in consecutive years (*i.e*., 2018-2020) in Little Turkey Lake and that reported (i) similar cyanobacterial community composition between sampling years, and (ii) variability in the relative abundance of cyanobacteria within the broader bacterial community (Cameron et al., 2022). Here, cyanobacteria were a statistically significant portion of the bacterial community in 2022, but not 2018, underscoring the potential for significant interannual variability in bacterial community structure and the need for further investigation into environmental drivers of these differences in oligotrophic systems.

Critically, the increased availability and rapidly decreasing costs of next-generation sequencing tools have eliminated most of the barriers to their wider use in resource management and the drinking water industry. The insights enabled by these approaches allow for proactive rather than reactive management of drinking water supplies (Chapman, 2010). Although amplicon sequencing has revolutionized our ability to study microbial communities, these data do not quantify abundance of community members. Rather, they only indicate community composition and relative abundance (Gloor et al., 2017, 2016) which introduces challenges in the interpretation of changes in community structure. For example, an observed decrease in relative abundances of cyanobacteria may only be an artifact of increased relative abundances of sequences classified to other taxonomic groups and not reflective of an actual decrease in cyanobacterial populations. Thus, to best inform cyanobacterial community dynamics for the purposes of resource and risk management, traditional cell enumeration techniques (e.g., flow cytometry; Patel et al., 2019) or other molecular approaches (e.g., qPCR; Chiu et al., 2017) that provide community density information are needed in conjunction with amplicon sequencing.

The sporadic diurnal variations in cyanobacterial populations and the differences between interconnected oligotrophic lakes that were reported herein emphasize the challenge in developing lake and reservoir monitoring protocols in absence of sufficient baseline knowledge of ecological community structures in the specific systems of interest. Here, cyanobacterial relative abundances fluctuated substantially between the hours of 9 a.m. and 4 p.m., demonstrating that large, generalized time frames are insufficient for characterizing cyanobacteria occurrence and dynamics in aquatic systems and associated threats to water quality and treatability. This work specifically demonstrated that following typical guidance that limits sample collection to a discrete depth in stratified lakes or a single timepoint may result in vast underestimation of cyanobacterial relative abundances. While cyanobacteria sampling protocols and guidelines do occasionally mention that time and depth of sample collection can affect outcomes, they are often vague and refer to collection of “surface waters” at times “later in the day” or from “10 a.m. to 3 p.m.” (Table S7), and do not address the potential for or implications of random non-diurnal migrations of smaller taxa. The present investigation fills a critical knowledge gap by reporting these insights for oligotrophic systems.

The design of sample collection schemes is closely linked to the utility of the information that is delivered using these approaches. The analysis provided herein has delivered clear evidence to demonstrate the need for integrating multi-timepoint, multi-depth discrete sampling guidance into lake and reservoir monitoring programs to describe cyanobacterial community dynamics and signal change to inform risk management associated with the potential for cyanotoxin production. This work further emphasized that cyanobacteria are present in oligotrophic lakes and their community structure varies (i) diurnally, (ii) across the depth of the water column, (iii) interannually within the same lake and (iv) between different lakes that are closely interconnected within the same watershed. Ignoring this variability and reducing sample numbers can lead to a false sense of security and missed opportunities to identify and mitigate changes in trophic status and associated risks such as toxin or taste and odor production, especially in sensitive, oligotrophic systems.

## 5. Conclusions

■ The potential for diurnal migration should be reflected in cyanobacterial monitoring programs through the inclusion of multiple-sampling times or conducting sampling at an ecologically significant time of day.
■ Sampling of lakes should not be restricted to the surface and should include discrete sampling and multiple depths to reflect spatial heterogeneity of cyanobacterial communities present in the water column.
■ Cyanobacterial communities are highly dynamic and may vary significantly in composition, prevalence and occurrence times between sampling years at the same sampling site.
■ Cyanobacterial monitoring may be enhanced through the incorporation of system characteristics (e.g., thermal stratification) and characterization of communities for design of sampling protocols that are system specific.

## Acknowledgements

We acknowledge the support of the *for*Water NSERC Network for Forested Drinking Water Source Protection Technologies [NETGP-494312-16]. We are also grateful for the continued support of Natural Resources Canada and Environment and Climate Change Canada in sample collection at Turkey Lakes Watershed Research Station.

