## Supplementary figures for "Sporadic Diurnal Fluctuations of Cyanobacterial Populations in Oligotrophic Temperate Systems Can Prevent Accurate Characterization of Change and Risk in Aquatic Systems"

**Figure S1** The Turkey Lakes Watershed location and lake sites. The watershed is located approximately 50 km north of Sault Ste Marie, Ontario as indicated by the marker on the map. The watershed consists of 4 interconnected basins visualized here using topographic maps generated through the Ontario Ministry of Natural Resources & Forestry.

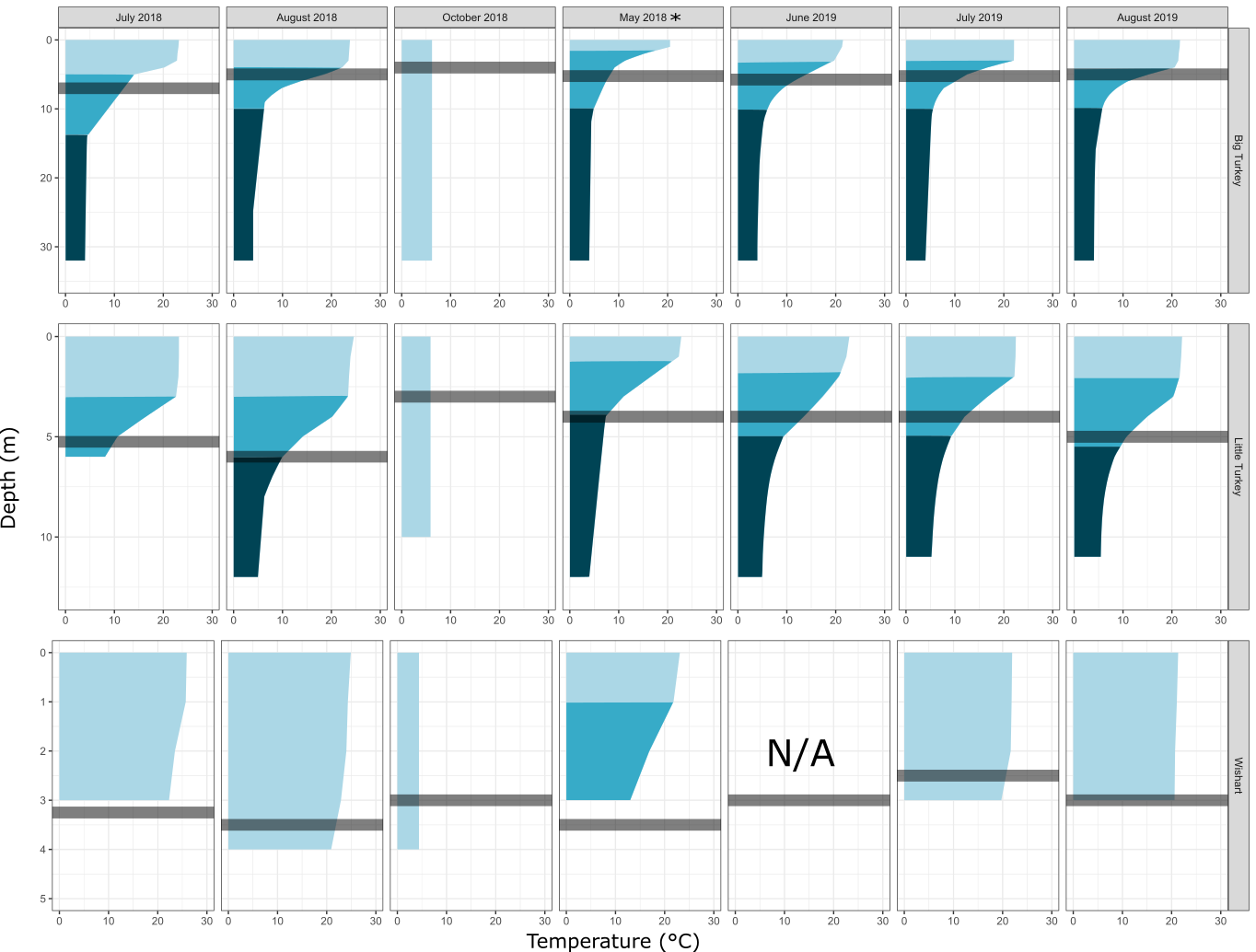

**Figure S2** Water column temperature profiles in ice-free months of 2018 and 2019. Water column temperature profiles were collected during ice-months in Big Turkey (Max depth = 37 m), Little Turkey (Max depth = 13 m) and Wishart Lake (Max depth = 4.5 m). Secchi depth, which is used as a sampling depth in the studies in this thesis, is indicated with the shaded grey bar. Thermally stratified layers are identified as epilimnion (light shade), metalimnion (medium shade) and hypolimnion (dark shade). Notably, water column profiles for June 2019 in Wishart Lake were not available. May 2019 data in all three lake sites was also unavailable but to demonstrate previous thermal stratification trends, data from May 2018 have been provided. Temperature profile data not available for 2022.

**Figure S3** Phylogenetic tree confirming the taxonomic affiliation of operational taxonomic units classified as cyanobacteria at the phylum level. Sequences were aligned to the Cyanoseq database and the tree was constructed using an unrestricted model in iqtree with 1000 boot straps and visualized using iTOL. Branches coloured in increasing intensity of blue represent high support (bootstrap > 0.8). Operational taxonomic units identified in this research are indicated with black boxes on the inner colour strip ring. Taxonomic lineages of clades where operational taxonomic units were placed were coloured according to family or order level.
